# Real-time sampling of travelers shows intestinal colonization by multidrug-resistant bacteria to be a dynamic process with multiple transient acquisitions

**DOI:** 10.1101/827915

**Authors:** Anu Kantele, Esther Kuenzli, Steven J Dunn, David AB Dance, Paul N Newton, V Davong, Sointu Mero, Sari H Pakkanen, Andreas Neumayr, Christoph Hatz, Ann Snaith, Teemu Kallonen, Jukka Corander, Alan McNally

**Affiliations:** Inflammation Centre, University of Helsinki and Helsinki University Hospital, Helsinki, Finland, POB 372, FI−00029 HUS; Human Microbiome Research Program, Faculty of Medicine, University of Helsinki, POB 21, FI-00014 University of Helsinki; Swiss Tropical and Public Health Institute, Basel, Switzerland; University of Basel, Basel, Switzerland; Epidemiology, Biostatistics and Prevention Institute, University of Zurich, Zurich, Switzerland; Institute of Microbiology and Infection, College of Medical and Dental Sciences, University of Birmingham, Birmingham, B15 2TT, UK; Lao-Oxford-Mahosot Hospital-Wellcome Trust Research Unit, Microbiology Laboratory, Mahosot Hospital, Rue Mahosot, Vientiane, Lao People’s Democratic Republic; Centre for Tropical Medicine and Global Health, Nuffield Department of Medicine, University of Oxford, Old Road Campus, Roosevelt Drive, Oxford, OX3 7LG, UK; Faculty of Infectious and Tropical Diseases, London School of Hygiene and Tropical Medicine, Keppel St, Bloomsbury, London WC1E 7HT, UK; Department of Infectious Diseases and Hospital Hygiene, Cantonal Hospital, St. Gallen, Switzerland; Institute of Basic Medical Sciences, Faculty of Medicine, University of Oslo, Oslo, Norway; Wellcome Sanger Institute, Cambridge, United Kingdom; Department of Mathematics and Statistics, University of Helsinki, Helsinki, Finland

**Keywords:** antimicrobial resistance, AMR, MDR, ESBL, whole genome sequencing, genomics, mcr, colistin, LMIC

## Abstract

**Background:** Antimicrobial resistance (AMR) is highly prevalent in low- and middle-income countries (LMIC). International travel contributes substantially to the global spread of intestinal multidrug-resistant gram-negative (MDR-GN) bacteria. Of the 100 million annual visitors to LMIC, 30–70% become colonized by MDR-GN bacteria. The phenomenon has been well documented, but since sampling has only been conducted after travelers’ return home, data on the actual colonization process are scarce. We aimed to characterize colonization dynamics by exploring stool samples abroad on a daily basis while visiting LMIC.

**Methods:** A group of 20 European volunteers visiting Lao People’s Democratic Republic for three weeks provided daily stool samples and filled in daily questionnaires. Acquisition of extended-spectrum beta-lactamase-producing gram-negative bacteria (ESBL-GN) was examined by selective stool cultures followed by whole-genome sequencing (WGS) of isolates.

**Results:** While colonization rates were 70% at the end of the study, daily sampling revealed that all participants had acquired ESBL-GN at some time point during their overseas stay, the colonization status varying day by day. WGS analysis ascribed the transient pattern of colonization to sequential acquisition of new strains, resulting in a loss of detectable colonization by the initial MDR-GN strains. All but one participant acquired multiple strains (n=2–7). Of the total of 83 unique strains identified (53 *E. coli*, 10 *Klebsiella*, 20 other ESBL-GN species), some were shared by as many as four subjects.

**Conclusions:** This is the first study to characterize in real time the dynamics of acquiring MDR-GN during travel. Our data show multiple transient colonization events indicative of constant microbial competition.

## Background

Antimicrobial resistance (AMR) poses a serious threat to human health worldwide [1]. The rapid global spread of multidrug-resistant (MDR) clones of *Escherichia coli, Klebsiella pneumoniae* and other Enterobacteriaceae raises an alarming public health concern [1]. Worldwide dissemination of successful clones such as *E. coli* ST131 has been the primary driver in extended-spectrum beta-lactamase (ESBL)-producing *E. coli* (ESBL-Ec) becoming prevalent among clinical isolates [2, 3]. Correspondingly, the global spread of carbapenemase-producing clones of *E. coli* such as ST410 [4] and ST167 [5], and *K. pneumoniae* clones such as CG258 and ST11 [6, 7] largely accounts for the rapid emergence of carbapenem resistance in clinical isolates of gram-negative pathogens worldwide.

The literature shows international travel to be strongly associated with acquisition of MDR-GN strains, mostly ESBL-*Ec* [8–15]. The carriage rate of intestinal MDR-GN bacteria is highest among inhabitants of the Indian subcontinent and Southeast Asia, followed by Africa and South America [16]. Not surprisingly, travelers visiting these high-risk regions are at substantial risk of acquiring MDR-GN bacteria [15]. Colonization occurs even during short visits [8–15] and without antimicrobial use [17] and can last for months or even over a year [8,12,13] and lead to further spread after return home [8, 13]. Genome-level analysis of MDR strains colonizing travelers shows that newly acquired MDR strains tend to displace resident intestinal commensal *E. coli* strains alongside new non-MDR strains, such that the pre-travel population remains as a minority [18].

In previous studies of travelers’ intestinal colonization by MDR-GN bacteria, samples have been taken immediately prior to travel and upon return to home country [8–14]. As such the dynamics of this competitive colonization process are unknown. Here we present a longitudinal study carried out with volunteers visiting the Lao People’s Democratic Republic (Laos). Combining daily samples from these travelers during their stay and fine-scale genomic analysis enabled us to characterize the colonization dynamics.

## Methods

### Study aim, design and setting

To obtain data on the dynamics of intestinal MDR bacteria acquisition, we characterized the colonization process by ESBL-GN by sampling participants’ stools on arrival, each day abroad, and at departure. The study was conducted over 22 days. The specimens were examined for presumptive ESBL-GN by culture, and the isolates were analyzed by whole-genome sequencing. The study protocol was approved by the Ethics Committee of the Helsinki University Hospital and the Ethikkommission Nordwest-und Zentralschweiz. All subjects provided written informed consent.

### Characteristics of volunteers, samples, and travel destination

Volunteers were recruited prospectively among participants at a medical course held 19 September – 9 October, 2015 in Vientiane, Laos. We also invited the recruitees’ companions to volunteer. While staying there, each volunteer was asked to provide daily stool samples. Those collected within the first two days were considered as baseline samples and the final stools as departure samples. Questionnaires assessing background information and travel-related data were used at recruitment and before departure. During the stay, the volunteers were asked to complete a health card to record gastrointestinal symptoms, food habits, and medication use each day. Travelers’ diarrhea (TD) was defined as passage of ≥ 3 loose or liquid stools per day. Any ESBL-GN not found in the baseline samples but detected in one or more stool samples taken later were defined as travel-acquired ESBL-GN. Only volunteers providing at least five daily samples were included in the final subject group (Fig. 1).

**Figure 1.**
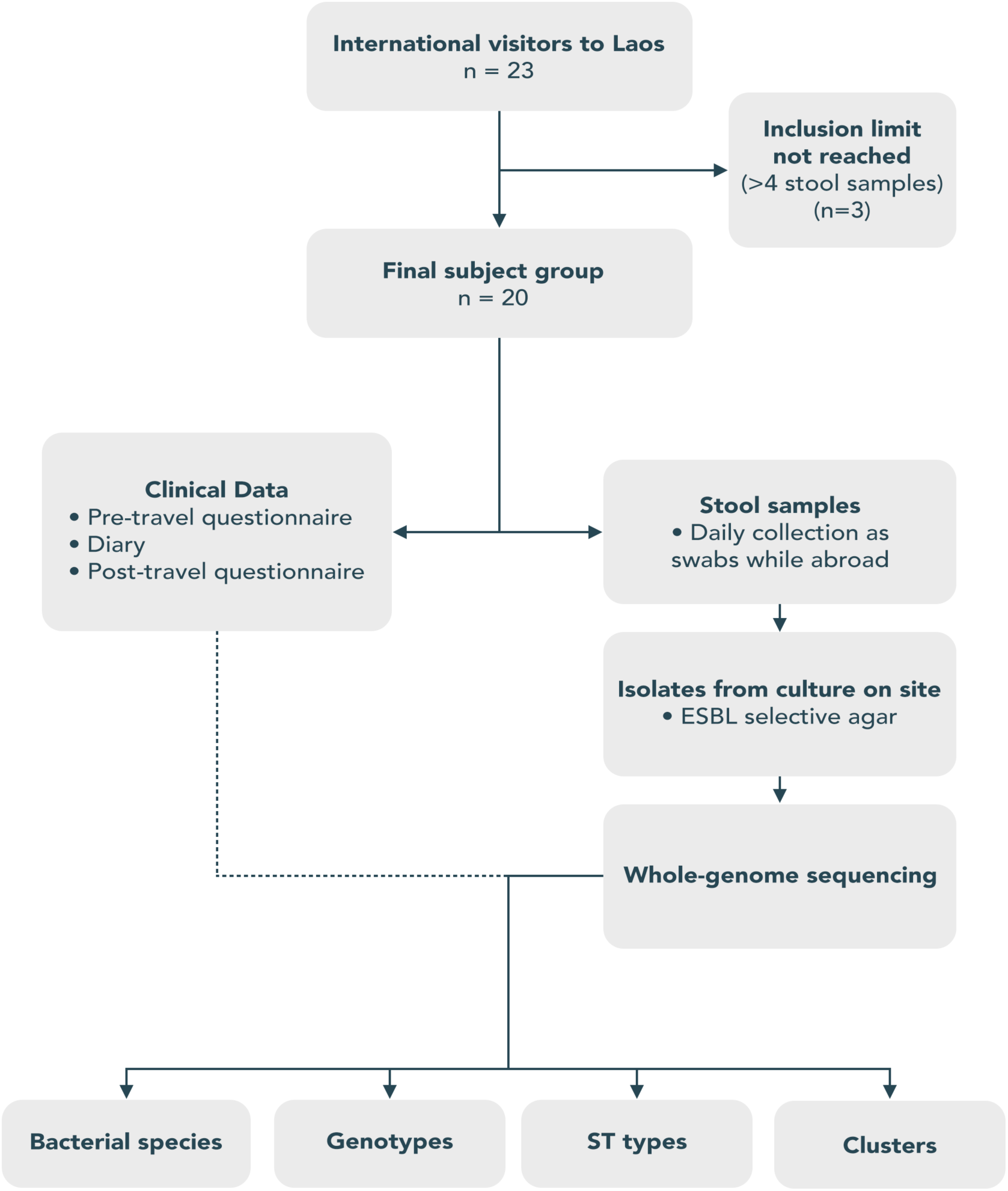
Overview of sampling and analysis workflow with inclusion criteria. This study recruited 23 participants who provided daily stool samples. These samples were screened for gram-negative extended-spectrum beta-lactamase-producing gram-negative (ESBL-GN) bacteria. Participants who produced less than four stool samples (due to constipation, for example) were excluded from the final study. Following selective culture, isolates were whole-genome sequenced and analysed to determine sequence type, resistance profiles, and genomic homology.

### Stool cultures and phenotypic susceptibility testing

The initial screening for presumptive ESBL-GN strains from stool samples was conducted in Vientiane at the Microbiology Laboratory of Mahosot Hospital by culture on CHROMagar™ ESBL agar plates (CHROMagar, Paris France). Phenotypically distinct colonies were subcultured and stored at −80°C in Protect tubes (Technical Service Consultants Ltd, Heywood, UK) along with the original swabs. The samples were then transported by plane on dry ice first to the Swiss Tropical and Public Health Institute in Basel, and then to the University of Helsinki (Finland) for analyses. In Finland, the bacteria were re-cultured on an equivalent chromogenic media, ChromID ESBL agar (BioMérieux, Marcy-l’Étoile, France), and a colony of all morphotypes present on the ChromID plate was subcultured and cryopreserved. The isolates were shipped on dry ice to the University of Birmingham for genome sequencing by the MicrobesNG facility (http://microbesng.uk). Libraries were prepared using the Nextera XT kit, and sequenced on the Illumina HiSeq platform.

### Genomic analyses

Illumina genome sequence reads were trimmed using Trimmomatic (V 0.3) [19] with a sliding window quality of Q15 and length of 20 base pairs. *De novo* assembled genomes were produced using SPAdes (V 3.13.0) [20]. Resulting assembled genomes were annotated using Prokka (V 1.11) [21]. Antibiotic resistance genes were detected in assembled and annotated genomes using Abricate (V 0.8.7, https://github.com/tseemann/abricate) and the Resfinder database. Prokka-annotated genomes were manually inspected to confirm the presence of resistance genes identified. MLST (V 2.15, https://github.com/tseemann/mlst) was used to verify species identification and assign classical sequence type designations to isolates. Where isolates were suspected of sharing recent source or transmission events, Snippy (V 4.3.6, https://github.com/tseemann/snippy) was used to map reads of isolates against the assembled genome of the earliest relative isolate. The number of SNPs between strains was determined using snp-dists (V 0.6.3, https://github.com/tseemann/snp-dists).

## Results

### Description of participants, travel, and symptoms during stay

A total of 23 volunteers were recruited, three of whom had to be later excluded for having only provided two samples (of note, ESBL-GN was found in them all). The final study population thus comprised 20 European volunteers. Of the volunteers, 50% were aged < 50 years, 55% were female, and 19 of them were medical doctors. The median age was 42.5 years (IQR 33.5−57.0), and the median duration of stay in Laos was 20 (IQR 12−21) days. Five participants (25%) had used antimicrobial medication during the previous year, three (15%) arrived directly from another tropical region, one (5%) had visited the tropics within the past three months and seven (35%) within the last year (Table S1). The group provided a total of 236 stool samples.

Over the sampling period, the volunteers stayed at three separate hotels, visited various restaurants either in small groups or all together, and participated in daily rounds at local hospitals. Four of the twenty participants reported TD, and one took antibiotics (Table S1).

### All participants were colonized by ESBL-GN, dominated by *E. coli*

ESBL-GN strains were detected in the primary samples of seven of the 20 volunteers, including three who had arrived directly from another Asian destination. For all volunteers at least one stool sample tested positive for ESBL-GN in culture by day 10 of their visit (Fig 2). Of the 236 fecal samples collected, 73.7% (n=174) contained detectable ESBL-GN, yielding a total of 306 isolates. *E. coli* was the most abundant species isolated, accounting for 219 of the 306 isolates, followed by *Citrobacter* spp. (28 isolates), *Klebsiella* spp. (16), *Acinetobacter* spp. (12), *Enterobacter cloacae* (11), and a number of other low-prevalence species including *Aeromonas* spp. and *Stenotrophomonas maltophilia* (Table S2). When allocating individual isolates to the participants, colonization by any given ESBL-GN during the study period was clearly transient in nature, with isolates detected in only one or a few samples obtained from any given individual, sometimes with days between isolation.

**Figure 2.**
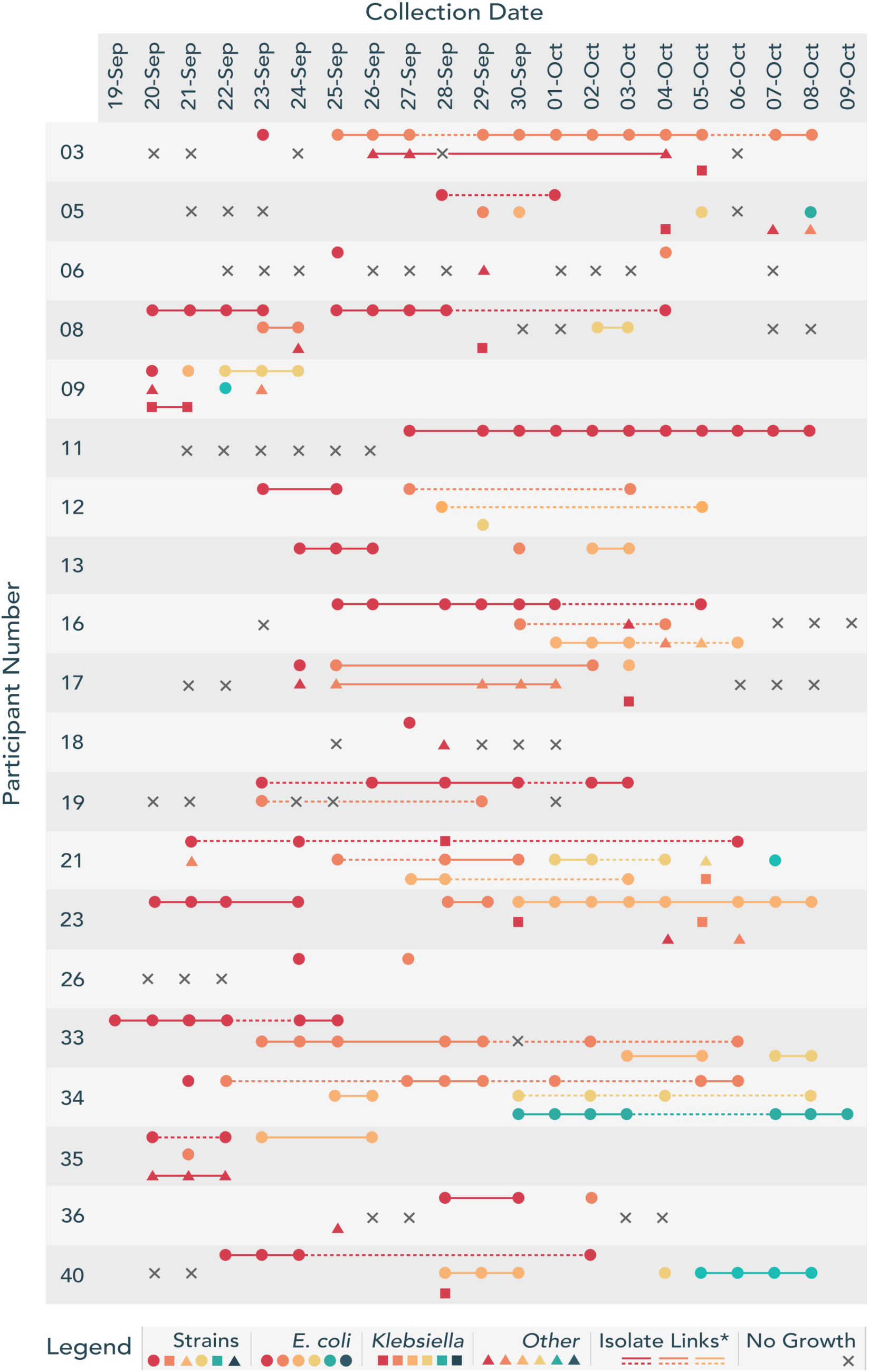
Colonization of participants by ESBL-producing gram-negative (ESBL-GN) over a 21-day period in Lao PDR. Patterns of colonization vary dramatically between participants. Some harbored a single dominant strain, whilst more transient and frequent strains were detected in the stool samples of others. Some participants appeared to have carried a considerably higher load of ESBL bacteria (e.g. 21, 34), while others only had one or two samples with ESBL isolates (e.g.18, 26). Solid lines represent uninterrupted and concurrent colonization by a single strain. Dashed lines represent maintenance of a single strain over multiple days interrupted by colonization by another strain of the same species or a period of no growth. Strains are shown by participant, with a maximum of five differing strains identified from the samples of a single individual. Due to the large number of constituent strains in the database, the color designations do not represent the same strains across multiple volunteers.

### Genomic analyses of isolates identified high variability in colonization both at strain and species level

The genomes of all 306 isolates were found to contain at least one antimicrobial resistance gene. The most prevalent ESBL gene type was *bla*_CTX-M_, found in a total of 226 isolates (74%), with CTX-M-55 (n=64), CTX-M-14 (n=58), CTX-M-159 (n=57), CTX-M-15 (n=30), and CTX-M-102 (n=25) as the most common types (Table S3). Some isolates contained multiple CTX-M types. Mobile colistin resistance genes (*mcr*) were found in 82 isolates (28%), all but two of which were *E. coli* (one *Aeromonas* sp., the other *K. pneumoniae*). Superimposing MLST designation (Figure 2, Figure S1) and *bla/mcr* gene type (Figure 3) onto each individual traveler’s isolates provided further insight into the transient nature of gut colonization. Whilst participant 11 was found to have contracted a single ST2067 strain of *E. coli* carrying CTX-M-15, the others were colonized by multiple STs of *E. coli* and co-colonized by other ESBL-GN species. For example, participant 34, who took azithromycin for TD on 21–23 September, was transiently colonized by five different *E. coli* strains over the study period, each with different *bla* gene repertoires. Participant 16 showed a regular flux between isolation of an ST38, ST93, and ST101 strain, whilst participant 40 had initial colonization by an ST48 strain later displaced by an ST38. All these strains have unique signatures of carriage of multiple *bla* genes, indicating that travelers are exposed to a large number of MDR-GN bacteria and MDR-conferring genes during the initial colonization process. All but one participant acquired multiple strains (n=2–7).

**Figure 3.**
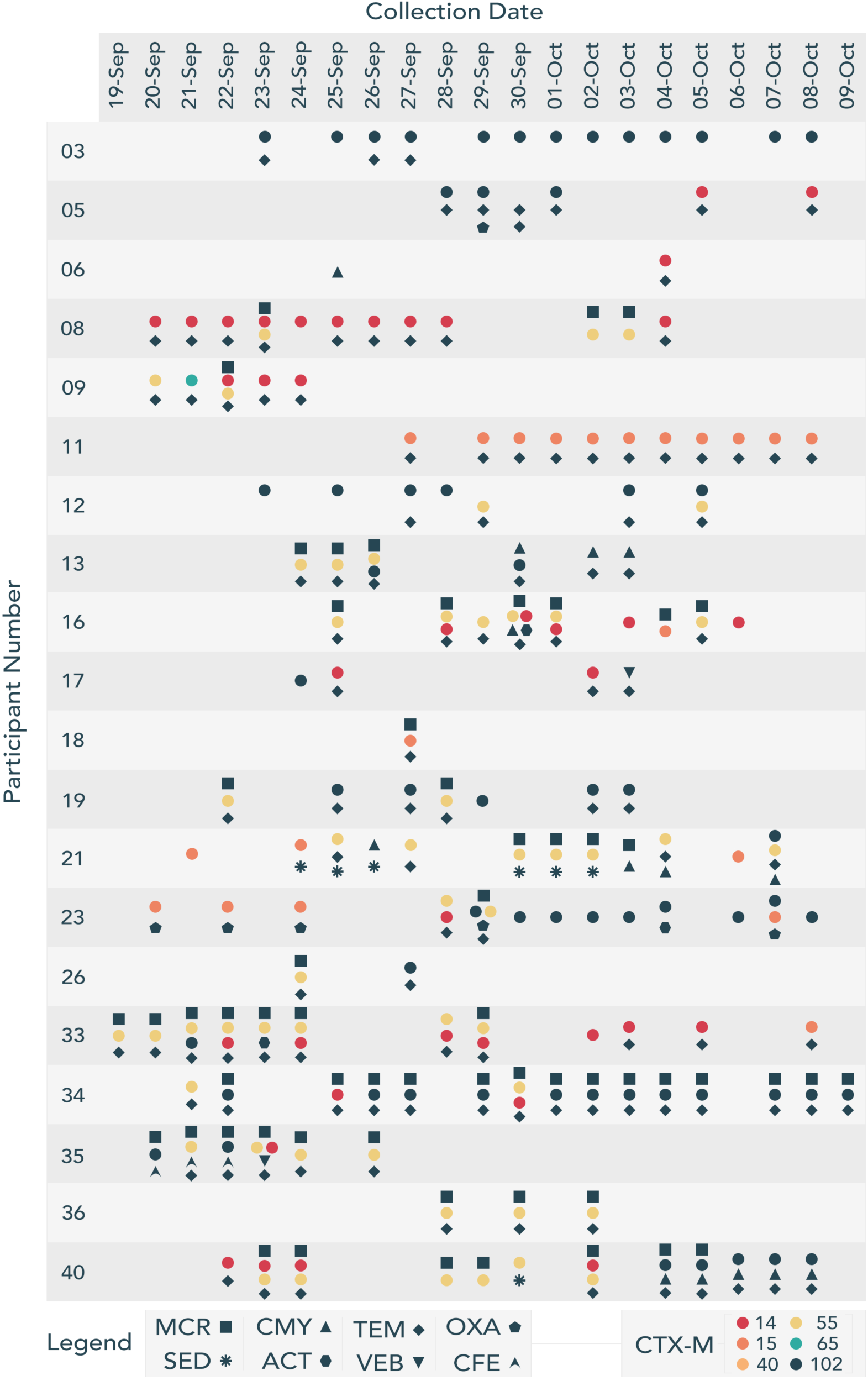
Resistance determinants identified in ESBL-GN isolates over a 21-day period in Lao PDR. We detected a highly diverse and abundant set of resistance determinants with numerous CTX-M variants, and a surprising abundance of MCR (28% carriage rate). We also found less common ESBL types, such as CMY, VEB, and CFE. Frequent patterns of resistance genes co-occurred in the same patient, overlapping with instances where a single colonizing strain was identified (detailed in Figure 2). Some of these also extended to other participants, (further characterized in Figure 5). Beta-lactamase genes are encoded using symbols, MCR = square, CMY = triangle, TEM = diamond, OXA = pentagon, SED = star, ACT = hexagon, VEB = inverted triangle, CFE = kite. CTX-M enzymes are encoded in colored circles, with CTX-M subtype encoded by different hues. CTX-M-14 = red, 15 = dark orange, 40 = pale orange, 55 = yellow, 65 = teal, and 102 = navy.

### Uncommon population structure of ESBL-GN isolates

We analyzed the population of *E. coli* isolates at MLST designation level (Figure 4). The most common *E. coli* sequence types identified were ST101, ST34, ST38, and ST195. These lineages are very uncommon in surveys of ESBL and carbapenem-resistant *E. coli*, both in Europe [3] and indeed in previous human isolates from Laos [22, 23]. Superimposing the presence of specific ESBL-Ec and *mcr* genes onto a phylogenetic tree of the *E. coli* isolates revealed *bla*_CTX-M_ genes to be ubiquitous throughout the sampled population of isolates, with *mcr* genes also widely distributed across the population (Figure S2). Analysis of the *K. pneumoniae* lineages showed isolates belonging predominantly to ST2176 and ST37, none of which are well characterized globally disseminated clones [6, 24].

**Figure 4.**
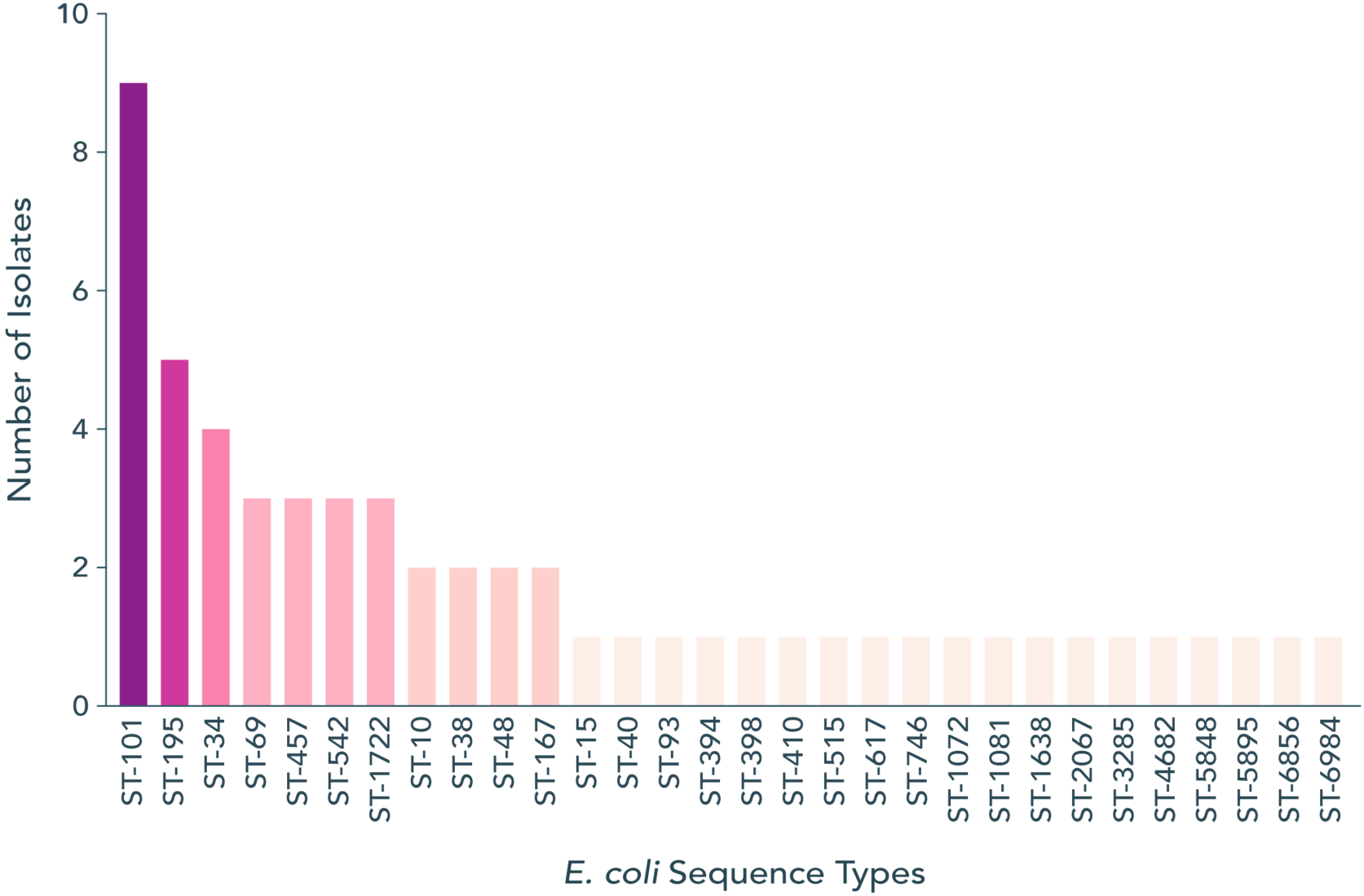
Abundance of unique sequence types amongst the traveler cohort over the 21-day study period. Isolates were examined using a high-resolution SNP analysis. Strains exhibiting genetic heterogeneity were considered unique, and their sequence type was recorded. ST-101 was the most abundant sequence type observed.

### Fine-scale genomic analysis identified common strains colonizing participants

When analyzing the population structure of the *E. coli* isolates, very little diversity was found in strains within each of the different lineages. To investigate relationships between isolates within the lineages, we carried out a high-resolution SNP analysis using the first isolated strain as a reference. Our data showed a number of common strains colonizing participants, often with zero SNPs’ difference between them (Figure 5). An identical ST515 strain colonized participants 6, 17, 33, and 5, whilst an identical ST38 colonized participants 5, 40, and 13. Participants 19 and 34 shared an identical ST34 strain, whilst participants 6 and 21 shared an identical ST385 strain. Participant 11 was colonized on day 8 by an ST2067 isolate that was also isolated from a companion of Participant 11 (11B) on the following day (Figure 5).

**Figure 5.**
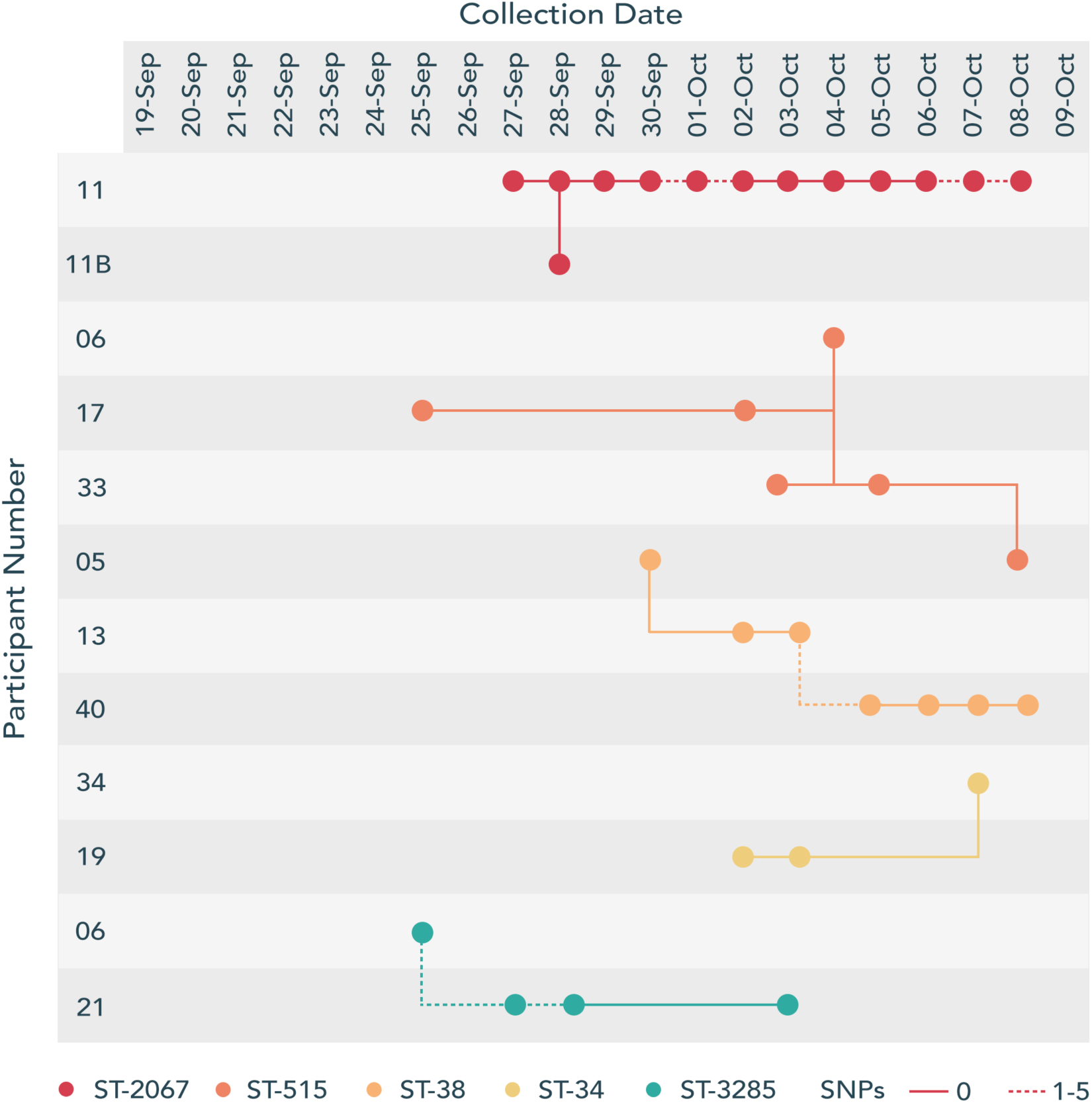
Linkage of isolates between participants. High resolution SNP analysis identified several instances of a single strain colonizing multiple participants. These strains differed by ≥5 SNPs, with the most prolifically shared strain (ST-515) found in participants 06, 17 and 33. Participant 11B represents a contact of participant 11, who was found to be colonised by a shared strain. Solid lines represent isolates that were identical (i.e. 0 SNP’s difference). Dashed lines represent isolates that were found to contain between 1-5 SNPs.

## Discussion

Combining personal data of 20 European visitors to Laos with fine-scale genomic analyses of the strains isolated from their daily fecal samples, we demonstrated that colonization by an ESBL-GN strain during travel to endemic regions is an extremely dynamic process. We identified a constant influx of newly acquired ESBL-GN strains in all but one of the twenty participants. Over the duration of their visits, the volunteers were colonized by up to seven different strains, and often acquired multiple ESBL-GN species. Few traveler studies have employed genome-level analyses [18], but several have reported isolating more than one new colonizing ESBL-GN strain from post-travel samples [10–14]. Our data reveal the true scale and complexity at which drug-resistant bacteria colonize the intestinal tract during travel, demonstrating that it has been seriously underestimated, both with respect to rates of volunteers acquiring MDR-GN and number of individual strains contracted. In addition, our data clearly show that several of our participants lost some of their travel-acquired ESBL-GN strains while still abroad. This indicates that previous studies solely employing pre- and post-travel sampling have under-reported the actual extent to which travelers are colonized by ESBL-GN. The result is especially relevant if we assume that MDR-GN simply drop below detection level but do not vanish completely, only to resurface once selection pressure increases again. There is a potential caveat to our study in that the apparent cyclic disappearance and re-appearance of strains may be related to the sensitivity of the culture methods used, and colonizing strains may occasionally have been missed when picking colonies for sequencing.

Our fine-scale genomic analysis enabled identification of a number of strains shared by our volunteers. Some of the strains colonized up to four participants, often with zero SNPs’ difference across the entire genome, and with a maximum of five SNPs’ difference between shared strains. Whilst direct transmission cannot be confirmed, the clonality of the isolates suggests that the two colonization events did not result from exposure to a common environmental reservoir. Such reservoirs are generally colonized by bacteria for extended periods of time, which leads to extensive diversity within the bacterial population [25, 26]. Thus, direct transmission or acquisition through common exposure such as consumption of food or water appears the most likely explanation.

The population of ESBL-Ec isolates in this study has an unexpected composition. Epidemiological surveys of multidrug-resistant *E. coli*, especially those focusing on ESBL strains, are dominated by *E. coli* ST131 [27]. Epidemiological investigations carried out on *E. coli* in Laos have also shown ST131 to be the dominant drug-resistant lineage in the country [22, 23]. However, we isolated no ST131 strains, and ST38 was the only lineage in our study that has been reported in studies previously conducted in Laos. These studies investigated both intestinal colonization isolates [23] and clinical blood stream isolates [22], suggesting that the ESBL-Ec population in Laos may be particularly dynamic and prone to frequent fluctuation. Interestingly, ST101, ST34, and ST195, and other lineages frequently isolated in our study have never been reported as clinical ESBL-Ec isolates in countries such as the United Kingdom where high-quality longitudinal data are available [3]. Some reports describe ESBL-Ec ST101 [28], but none report ST34 or ST195. *E. coli* ST38 has been extensively reported as an ESBL-Ec strain isolated both from humans and animals [29–31].

Most previous studies among travelers have only described acquisition of ESBL-*E. coli.* A few have reported multiple findings of ESBL-DEC (diarrheagenic *E. coli*) [32] or single findings of ESBL-*Klebsiella* [11, 14] or carbapenemase-producing *E. coli* [12, 13]. Our data show, in addition to ESBL-*E. coli* (219 of 306 strains, 72%) a substantial number of ESBL-producing non-*E. coli* Gram-negative bacteria, such as *Citrobacter* (9%), *Klebsiella* (5%), *Acinetobacter* (4%), and *Enterobacter cloacae* (4%) and even low numbers of *Aeromonas spp.* and *Stenotrophomonas maltophilia*. This finding may be ascribed to an especially high rate of exposure to a variety of MDR-GN bacteria, since 19 of our volunteers attended a course of tropical medicine which included daily clinical rounds at local hospitals. High MDR-GN colonization rates have been reported among travelers hospitalized in the tropics [33].

The complete absence of carbapenemase-producing *E. coli* in these samples is also noteworthy. We screened for ESBL-producing strains, but carbapenemase producers would also have grown on the selective plates used. However, WGS analysis showed a complete absence of carbapenemase genes, a finding somewhat surprising in Southeast Asia where the prevalence of carbapenem-resistant Enterobacteriaceae is increasing [34]. Such isolates have only recently been reported in Laos [35], suggesting that carbapenem resistance has not yet become a major problem in the country. Additionally, other traveler studies have shown a low carriage rate of carbapenemase genes [36].

Even more striking were the extremely high levels of the mobile colistin resistance gene *mcr* in our *E. coli* isolates. The ST101 lineage which dominated our isolate collection has been identified as a driving lineage in the emergence of ESBL-Ec, and *mcr* positive *E. coli* in the region [37], but we observed the *mcr* gene across a large number of lineages, a finding which may be of substantial importance. This confirms earlier reports showing that travel exacerbates the global spread of not only ESBL genes but also *E. coli* strains carrying mobile colistin resistance genes [38].

## Conclusions

By combining real-time sampling of travelers with genome-level analyses, we have demonstrated that colonization by ESBL-GN during travel is an extremely dynamic process characterized by competition between resistant strains and an individual’s own microbiota. We have also shown that prevalent strains can colonize multiple travelers via shared routes such as transmission or acquisition from a common source. The challenge now lies in unravelling the mechanisms that underlie this process and competition between the clones, as well as finding tools to prevent colonization already at its initial stages.

## Declarations

### Ethics approval and consent to participate

The study protocol was approved by the Ethics Committee of the Helsinki University Hospital and the Ethikkommission Nordwest-und Zentralschweiz. All subjects provided written informed consent.

### Consent for publication

Not applicable.

### Availability of data and materials

Raw sequence data for all isolates is available via NCBI under the Bioproject accession number PRJNA558187.

### Competing interests

A. K. has received honoraria from Valneva and Immuron, and investigator-initiated grants from Pfizer and Valneva, none of them relevant to the submitted work.

All other authors report no potential conflicts of interest.

### Funding

AK was supported by the Finnish Governmental Subsidy for Health Science Research; the Scandinavian Society for Antimicrobial Chemotherapy Foundation; the Sigrid Jusélius Foundation; and the Finnish Cultural Foundation. SD and AM were funded by BBSRC [BBR0062611]. AS was funded by Wellcome [108876B15Z], and AM was also funded by MRC [MRS0136601] and the Royal Society [NA150363]. T.K. and J.C. were supported by JPIAMR grant Spark from Norwegian Research Council and J.C. was also supported by ERC [742158].

The funding sources had no involvement in study design, data collection, analysis, interpretation of data, writing of the report, and decision to submit the manuscript for publication

### Authors’ contributions

Authors AK, EK and SD contributed equally to this manuscript. Authors JC and AM contributed equally to this manuscript. Study concept, design and organization AK, EK; sample and epidemiological data collection: AK, EK, DABD, PNP, VD, SM, SHP, AN, CH; funding for WGS: AM, JC; genomic analyses: SD, AS, TK, JC, AM; data interpretation: SD, AS, JC, AM; data visualisation: SD, AS; drafting of manuscript AK, EK, SD, TK, JC, AM; critical comments on the manuscript: DABD, PNP, VD, SM, SHP, AN, CH, AS; all authors read and approved the final manuscript.

## Abbreviations

AMR: Antimicrobial resistance
DEC: diarrheagenic *Escherichia coli*
ESBL: extended-spectrum beta-lactamase
ESBL-Ec: extended-spectrum beta-lactamase (ESBL)-producing *E. coli*
ESBL-GN: extended-spectrum beta-lactamase-producing gram-negative bacteria
ESBL-PE: extended-spectrum beta-lactamase-producing *Enterobacteriaceae*
IQR: inter quartile range
Laos: Lao People’s Democratic Republic People’s Democratic Republic
LMIC: low- and middle-income countries
mcr: mobile colistin resistance genes
MDR: multidrug-resistant
MDR-GN: multidrug-resistant gram-negative bacteria
TD: travelers’ diarrhea
WGS: whole-genome sequencing

## Acknowledgements

We are very grateful to the volunteers and the staff of the Microbiology Laboratory, Mahosot Hospital who processed the samples. We are also grateful to Bounthaphany Bounxouei, past Director of Mahosot Hospital; to Bounnack Saysanasongkham, Director of Department of Health Care, Ministry of Health; to H.E. Bounkong Syhavong, Minister of Health, Lao People’s Democratic Republic for their support of the work of LOMWRU, which is funded by the Wellcome Trust.

