## Supplementary Materials for "Real-time sampling of travelers shows intestinal colonization by multidrug-resistant bacteria to be a dynamic process with multiple transient acquisitions"

Table S1. Demographics of participants in our study exploring acquisition of extended-spectrum beta-lactamase-producing E. coli by daily stool sampling over their visit to Lao People’s Democratic Republic in September–October, 2015.

| **ID** | **Age** (yrs) | **Sex** | **Country of origin** | **Arriving from** (if not country of origin) | **Date of arrival** | **Departure date** | **Travelers’ diarrhoea** | **Antibiotic use** |
| --- | --- | --- | --- | --- | --- | --- | --- | --- |
| 3 | 33 | male | Germany |  | 20 Sep | 10 Oct | 4 Oct |  |
| 5 | 67 | male | Switzerland |  | 20 Sep | 09 Oct |  |  |
| 6 | 52 | male | Finland |  | 20 Sep | 10 Oct | 28 Sep |  |
| 8 | 29 | female | Switzerland |  | 20 Sep | 09 Oct |  |  |
| 9 | 30 | female | Austria | Vietnam | 20 Sep | 26 Sept |  |  |
| 11 | 61 | female | Finland |  | 20 Sep | 10 Oct | 25/26 Sep |  |
| 12 | 64 | male | Austria |  | 10 Sep | 09 Oct |  |  |
| 13 | 38 | female | Switzerland |  | 20 Sep | 10 Oct |  |  |
| 16 | 62 | female | Switzerland |  | 19 Sep | 17 Oct |  |  |
| 17 | 53 | female | Finland | USA | 21 Sep | 10 Oct |  |  |
| 18 | 46 | male | Germany |  | 25 Sep | 04 Oct |  |  |
| 19 | 34 | female | Netherlands |  | 20 Sep | 03 Oct |  |  |
| 21 | 53 | female | Norway |  | 20 Sep | 07 Oct |  |  |
| 23 | 32 | male | Austria | Vietnam | 20 Sep | 09 Oct |  |  |
| 26 | 20 | female | Switzerland |  | 19 Sep | 28 Sep |  |  |
| 33 | 39 | male | Germany |  | 13 Sep | 24 Oct |  |  |
| 34 | 35 | male | Switzerland | Thailand | 20 Sep | 10 Oct | 20 Sep–7 Oct | 21–23 Sep |
| 35 | 63 | male | Switzerland |  | 19 Sep | 28 Sep |  |  |
| 36 | 52 | female | Germany |  | 25 Sep | 04 Oct |  |  |
| 40 | 37 | female | Germany |  | 19 Sep | 14 Oct |  |  |

Table S2 – Frequencies of ESBL positive gram negative taxa observed in the dataset. *E. coli* and other Enterobacteriaceae (e.g. *Citrobacter, Klebsiella*) were the most common species observed. Some genera were isolated in very low numbers, (e.g. *Stenotrophomonas, Aeromonas).*

| Taxa | Number of Isolates |
| --- | --- |
| *E. coli* | 219 |
| *Citrobacter* | 28 |
| *Klebsiella* | 16 |
| *Enterobacter cloacae* | 11 |
| *Acinetobacter* | 12 |
| Other | 20 |
| Total | 306 |

Figure S1 – Distribution of observed *E. coli* sequence types sorted by participant number and date. In some instances, there are clear single sequence types that longitudinally colonise a single participant (e.g. 1722, Participant 03). Other participants exhibit transient colonisation by multiple sequence types (e.g. Participant 33).

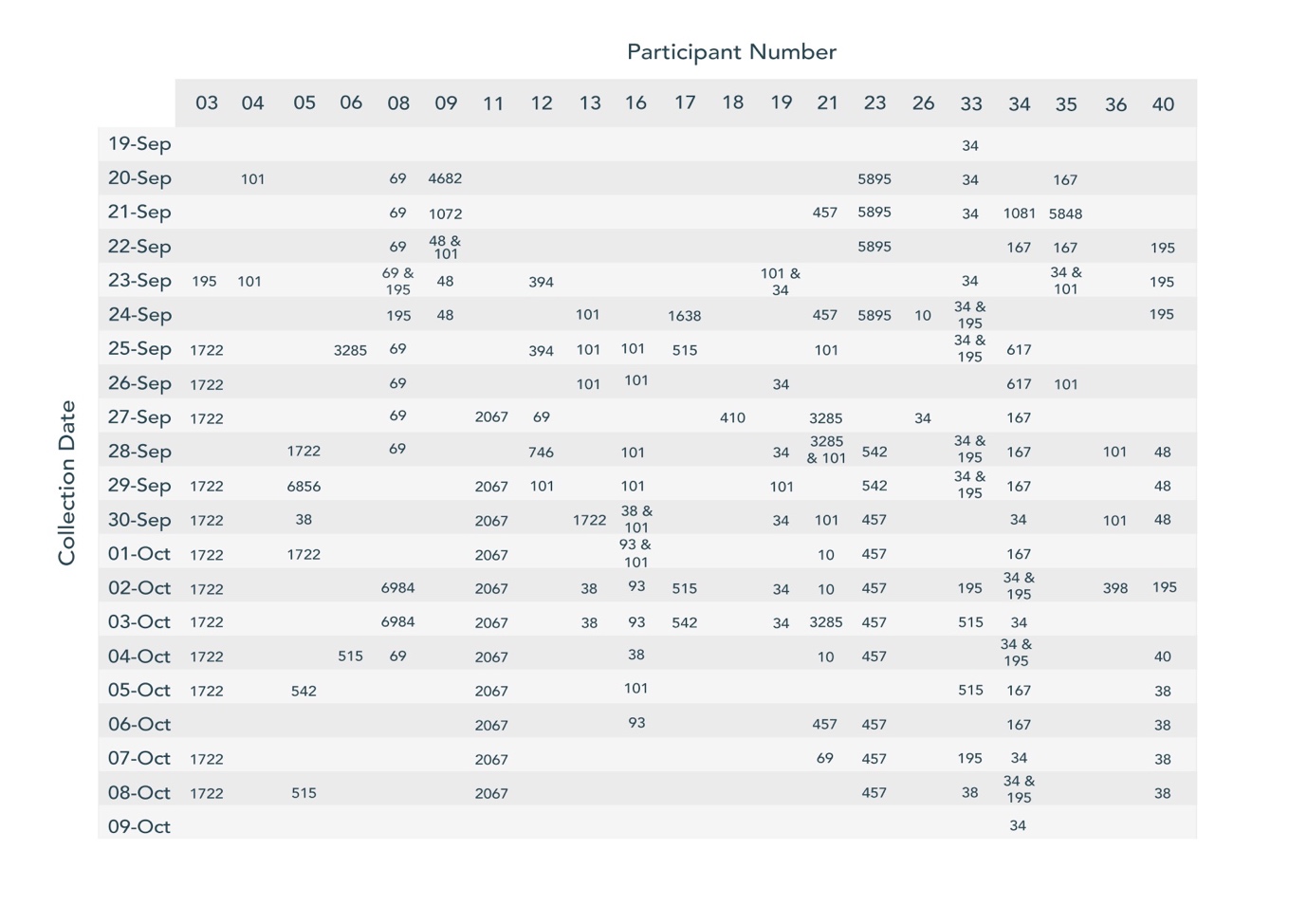

Figure S2 – Phylogeny of *E. coli* isolates shown with presence of observed beta-lactamase genes. The majority of isolates carried at least one type of CTX-M, and an alarming amount of isolates also carried colistin resistance gene MCR. Some less common beta-lactamase genes were also observed (e.g. ACT, ADC). Purple = gene present, orange = gene absent.

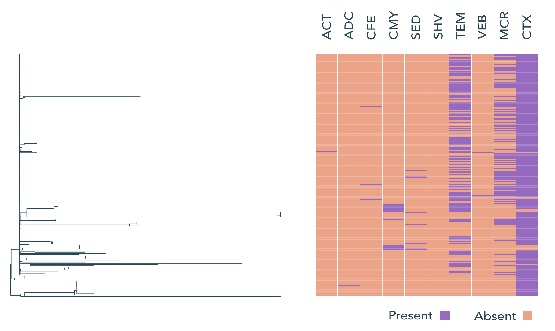

Table S3 – Total frequencies at which CTX-M subtypes were observed amongst the dataset. CTX-M-55 was the most common type observed, with other CTX-M types that are typically more dominant in other parts of the world (e.g. CTX-M-15) found to be less abundant.

| CTX-M Type | Number of Isolates with CTX-M Present |
| --- | --- |
| CTX-M-55 | 64 |
| CTX-M-14 | 58 |
| CTX-M-159 | 57 |
| CTX-M-15 | 30 |
| CTX-M-102 | 25 |
| CTX-M-40 | 2 |
| CTX-M-63 | 2 |
| CTX-M-164 | 1 |
| CTX-M-181 | 1 |
| CTX-M-196 | 1 |
| CTX-M-32 | 1 |
| CTX-M-65 | 1 |
| CTX-M-76 | 1 |
| CTX-M-77 | 1 |
